# Increased adaptability to rapid environmental change can more than make up for the two-fold cost of males

**DOI:** 10.1101/340927

**Authors:** Caroline M. Holmes, Ilya Nemenman, Daniel B. Weissman

**Affiliations:** Departments of Physics and Biology, Initiative in Theory and Modeling of Living Systems, Emory University*, Atlanta, GA 30322,* USA

## Abstract

The famous “two-fold cost of sex” is really the cost of anisogamy – why should females mate with males who do not contribute resources to offspring, rather than isogamous partners who contribute equally? In typical anisogamous populations, a single very fit male can have an enormous number of offspring, far larger than is possible for any female or isogamous individual. If the sexual selection on males aligns with the natural selection on females, anisogamy thus allows much more rapid adaptation via super-successful males. We show via simulations that this effect can be sufficient to overcome the two-fold cost and maintain anisogamy against isogamy in populations adapting to environmental change. The key quantity is the variance in male fitness – if this exceeds what is possible in an isogamous population, anisogamous populations can win out in direct competition by adapting faster.

## 1. Introduction

In most sexually reproducing species, the different sexes contribute different amounts of resources to offspring [1]. One fundamental way that they do this is via *anisogamy*: producing gametes of different sizes, so that one sex contributes more resources to the zygote. This anisogamy is in fact what defines the sexes, with females typically defined as the sex that invests more resources in its offspring [1]. Many sexual species take this asymmetry much further, with females providing essentially all the resources for offspring and males providing virtually nothing besides half of the offspring’s genome. In this case, assuming that male and female offspring require equal resources to reproduce, males impose their famous “two-fold cost” on females – parthenogenetic females could pass on twice as much of their genetic material as those who mate with males, since they would have the same number of offspring while being responsible for all of their genetic material.

While parthogenetic lineages have a short-term advantage, they essentially lose the ability to recombine, which can be crucial for generating variability that can be selected over longer time scales [2, 3, 4, 5, 6, 7, 8, 9, 10, 11]. However, this fact by itself does not explain the prevalence of males, as isogamous species (or, more generally, those in which both mating types invest equally in offspring, as in, for example, yeast [12]) retain all the benefits of recombination while still potentially producing twice as many offspring as anisogamous ones [13]. Why then are almost no multicellular sexual species isogamous, given that the fitness cost of anisogamy is so high?

The primary class of explanations for the prevalence of anisogamy is based on direct selection on the size of gametes and zygotes [14, 15, 16, 7, 17, 18, 19, 1]. For example, for widely-separated plants to produce large seeds, it will generally be much easier for a pollen grain to travel from one plant to another than for a half-seed (or for two half-seeds to meet somewhere in the middle). Related arguments also provide reasons why there are often exactly two mating types [20].

Another class of explanations for anisogamy, which we will focus on here, considers the influence of anisogamy on evolution. The key process at work is *sexual selection*, which we will use to refer to the fitness components relating to mating success. Sexual selection can also exist in isogamous species – for example, yeast produce and follow pheromone gradients to find mates [21]. As far as sexual selection acts only to produce assortative mating, in which high-fitness individuals preferentially mate with other high-fitness individuals, there is no necessary advantage for anisogamy, as isogamous populations can do this as well. However, because males in anisogamous populations can produce large numbers of sperm with little cost, they can experience an additional form of sexual selection, in which some males reproduce many times while others do not reproduce at all. While this is also possible to a limited extent in isogamous populations, having individuals that do not reproduce necessarily reduces the resources available for the next generation. By contrast, under anisogamy, if the males favored by sexual selection are the ones carrying “good genes” that increase other fitness components, this form of sexual selection can greatly enhance natural selection without a reduction in the reproductive output of the population [3, 22, 23], although it is unclear how often it actually does so in nature (see, e.g., [24]).

“Good genes” sexual selection allows anisogamous populations to greatly reduce their mutational load and the probability that deleterious mutations fix, potentially overcoming the two-fold cost if deleterious mutation rates are large [25, 26, 27, 28, 29, 30, 31, 32]. In other words, males can act as dead ends, where deleterious mutations go to die. Sexual selection can also help anisogamous populations adapt by increasing the fixation probability of beneficial mutations [26], but the magnitude of this effect is likely to be limited, as interference among mutations generally prevents their rate of incorporation from reaching levels that would balance the two-fold cost [33]. However, this interference limit does not apply to non-equilibrium selection on standing variation; in this case, even if all beneficial alleles start at frequencies such that they are essentially certain to be fixed, sexual selection can provide an advantage by allowing them to be fixed more rapidly [13, 24, 34]. The importance of this effect has long been known in animal breeders, who generally use extreme selection on males to rapidly improve stocks, with the thoroughbred stud Storm Cat, for instance, fathering over 1000 foals [35]. However, only one previous study has quantitatively considered how it might provide an advantage for anisogamy: Lorch et al. [36] observed that in simulated populations, sexual selection could produce a spike in the rate of adaptation to environmental change, although the observed advantage was not large enough to balance the two-fold cost. Here we use simulations of direct competition between isogamous and anisogamous populations to show that the increase in the rate of adaptation to environmental change can, in fact, be large enough to single-handedly balance the two-fold cost of males, and we quantify the conditions for it do so under a minimal model of sexual selection.

## 2. Model

We investigate the fitness effects of anisogamy by numerically simulating competition between a sexual anisogamous population and an equivalent sexual but isogamous population adapting to a sudden environmental change. We expect that the isogamous population will initially outcompete the anisogamous one because of the two-fold cost of males. However, we also expect that sexual selection will allow the anisogamous population to adapt faster, as the fittest males will produce very large numbers of offspring. If this increase in speed is large enough, the anisogamous population will overcome the two-fold cost before it goes extinct. We use simulations and approximate calculations to determine the parameter values for which this happens.

In our model, the isogamous population has two mating types; although they invest equally in offspring, we will refer to them as “females” and “males” since they play slightly different roles in the simulations. The total population size of all individuals together is fixed at *N* diploid individuals. Each generation, each anisogamous female produces *n eggs*, while each isogamous individual produces 2*n* gametes, corresponding to the two-fold cost of males. We emphasize that this model accounts for the general asymmetry in the parental investment in offspring, and not just in the gamete size. Each anisogamous male produces an effectively infinite number of sperm. In the first stage of selection, females (including isogamous “females”) compete with each other to place their gametes in the next generation, with each gamete being selected with probability proportional to the fitness of its mother. In the second stage of selection, males compete with each other to fertilize the successful female gametes. Anisogamous males only fertilize anisogamous female eggs and isogamous “males” only fertilize anisogamous “female” gametes, so there is no competition between mating systems in this stage and no interbreeding between the mating systems. That is, the competition between the isogamous and the anisogamous populations only occurs when selecting the female gametes that will mate and contribute to the following generation. Male gametes are selected with probability proportional to the fitness of their father. To keep the sex ratio fixed, each mating produces exactly two offspring, which are genetically identical except that one is male and one is female. We do not expect this constraint to significantly affect our conclusions.

Each individual is diploid, with a genome consisting of *L* loci. All loci are unlinked, i.e., gametes sample one of the two parental alleles at each locus independently. Each locus is binary, with allele 1 conferring an advantage in log fitness *s* over allele 0, with no epistasis. Thus if an individual has genotype X, with *X_k_* ∈ {0, 1, 2} being the number of 1 alleles that the individual has at locus *k*, its fitness *w* is:

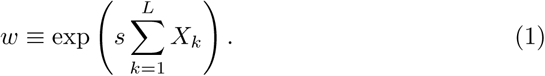

Technically, this is the individual’s breeding value for fitness rather than fitness itself (defined as the expected number of offspring), which also depends on sex and mating system. Explicitly, while an isogamous individual cannot have more than 2*n* offspring and an anisogamous female cannot more than *n* offspring, regardless of their value of *w*, an anisogamous male with extremely high w could, in principle, sire all of the anisogamous offspring in the next generation.

We are modeling a situation in which an environmental shift has just changed the selection on standing variation. For simplicity, we assume that all the alleles were previously neutral, with starting frequencies *F_k_*(*t* = 0) drawn independently from the distribution ([37], Eq. (9.3.3)):
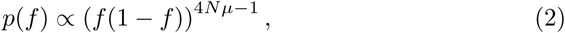

where *µ* is the mutation rate per locus per generation. At the beginning of the simulation, each individual’s alleles are drawn independently at each locus according to their frequency *F_k_*(0), i.e., the population is in linkage equilibrium, up to stochastic effects.

### 2.1. Simulated parameter values

For the genome length *L*, we considered between 10 and 1000 loci. The selective advantage *s* of each allele ranged between 0 and 0.5, with each simulation run having a single value for all loci. The loci and alleles in these simulations should be understood as linkage blocks – the longest stretches of genome that can be treated as effectively unlinked – rather than individual nucleotides. For instance, *L* = 100 and *s* = 0.1 might correspond to a human-sized genome of 3 gigabases, viewed as being composed of “loci” of 30 megabases each, each potentially contributing variance in log fitness of 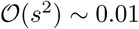. The population size was *N* ∈ [100, 10000]. Each simulation started with an equal number *N*/2 of anisogamous and isogamous adults and continued until one mating system drove the other to extinction. Since the effects described here are largely consistent across population sizes varying over orders of magnitude, as we have verified numerically, for concreteness, we use *N* = 1000 for all figures. We used mutation rate *µ* = 0.01 in (2), but did not actually include new mutations in the course of our simulations, as their effect is expected to be negligible given the large amount of standing variation and the short time needed for one population to out compete the other – e.g., < 10 generations in Fig. 1. Like *s*, *µ* should be understood as an effective parameter describing a linkage block rather than an individual nucleotide. As long as *Nμ* ≳ 1 (as it always is in our simulations), each linkage block will begin with standing variation and will harbor variance in log fitness on the order of the maximum value of ~ *s*^2^. As we are concerned with the effect of anisogamy on the maximum possible rate of adaptation, the simulated parameter values correspond to extremely strong selection, much stronger than is typically observed in natural populations; we consider the relationship with natural dynamics in the Discussion. For every plotted parameter combination, we ran 100 independent simulations to calculate averages, which ensured that statistical fluctuations are much smaller than the means.

**Figure 1:**
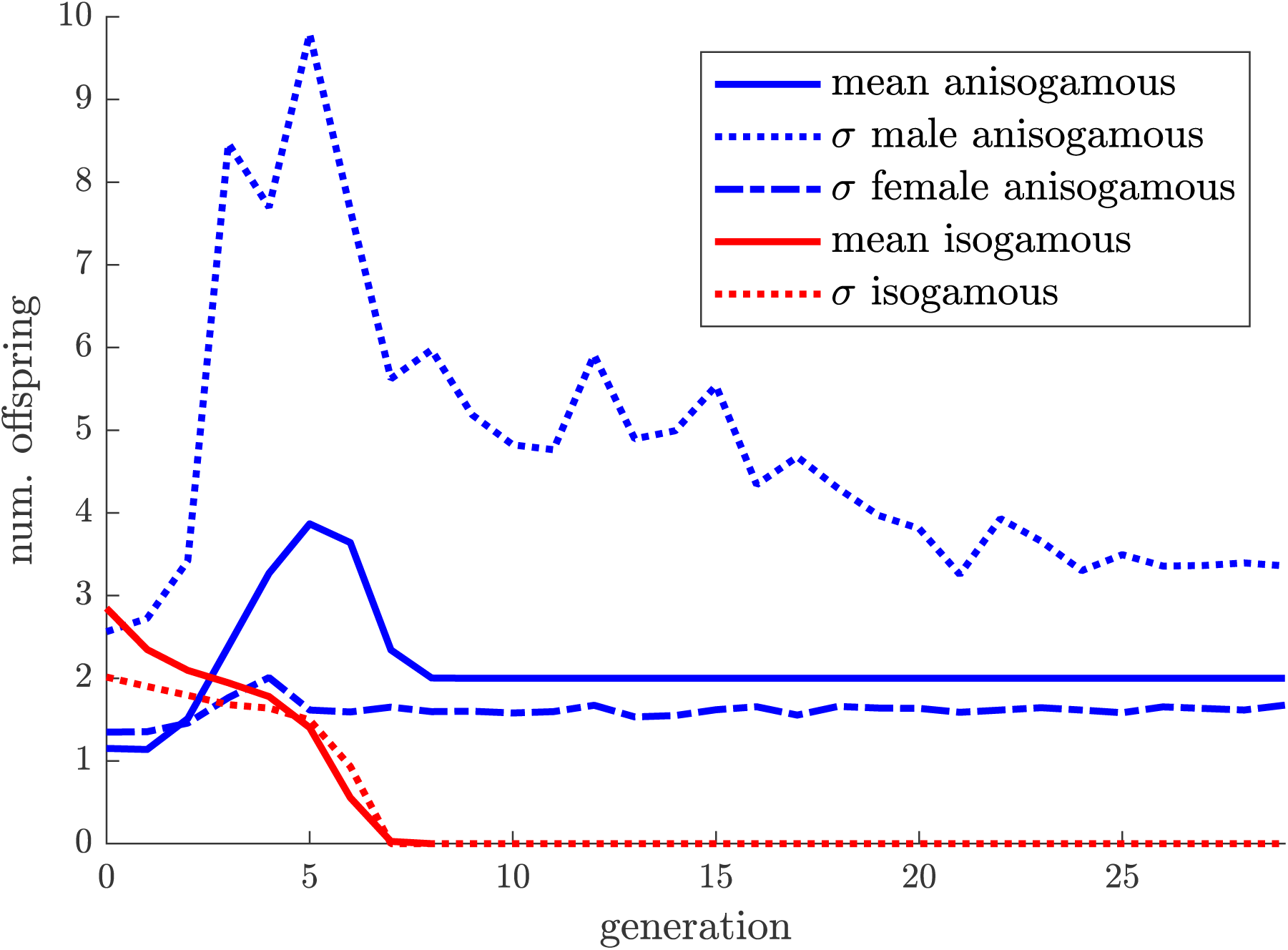
Trajectory of a single typical simulation run, showing the mean and standard deviation of the number of offspring per individual over time for isogamous and anisogamous individuals. In the first generation, isogamous individuals produce more than twice as many offspring as isogamous individuals due to the two-fold cost of males and stochastic effects, but the isogamous population adapts faster and drives them extinct by generation 7. The increased rate of adaptation is driven by the large spike in the variance in offspring number among males around generation 5, which cannot be matched by the isogamous population because all individuals have upper limits to their reproductive capacity. Parameters: *N* = 1000, *n* = 3 eggs per anisogamous female, *L* = 300 loci, and *s* = 0.3.

## 3. Results

Fig. 1 demonstrates dynamics of the realized fitness (number of offspring) of mating systems in a single typical run of the simulation, where anisogamy outcompetes isogamy. The mean fitness of the isogamous individuals starts at more than twice that of the anisogamous ones, because of the two-fold cost of males and stochasticity in the distribution of alleles and in reproduction (note that the mean fitnesses of anisogamous males and anisogamous females are the same since each offspring has one father and one mother). However, the mean anisogamous fitness then increases rapidly, eventually winning the competition before settling down at 2, the replacement rate. The figure also shows that the spike in the mean realized fitness of the anisogamous individuals coincides with a dramatic spike in the standard deviation of the realized fitness distribution of males: with a mean of ~ 3 offspring, the standard deviation of offspring per male reaches 10, so that some males can have 20 or more offspring, while others have zero. At the same time, as expected, the standard deviation of the realized fitness distribution of females barely increases (and always stays significantly below that of the males). This illustrates that the realized fitness of females is limited since the maximum offspring number is not more than the reproductive capacity *n* = 3, and there is, therefore, room to procreate even for not very fit females. In addition, this verifies our intuition that it is the variance in the number of offspring for males that drives the fitness increase.

Fisher’s fundamental theorem of natural selection states that the rate of increase of mean fitness is equal to the variance in fitness. Thus the rate of adaptation is limited to 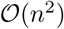 under isogamy, while anisogamous populations can adapt faster via selection on males, and we predict that this difference will affect the success of the anisogamous population once the variance in breeding values for fitness *w* approaches 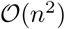, or, equivalently, log var(*w*) ~ log *n*. The populations start at linkage equilibrium and, since all loci are unlinked, we expect them to remain close to it. The fitness breeding values will therefore be approximately log-normally distributed within populations, with log fitness having variance 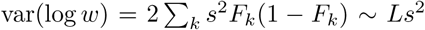, since each locus is an independent binomial variable. Since *w* is log-normal, the log of its variance is approximately 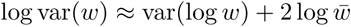, where 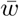 is its mean. Near the replacement-level mean fitness of 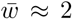 offspring, we therefore roughly need var(log *w*) ~ log *n* for the fit isogamous individuals to be hitting the limit of their reproductive capacity, or *Ls*^2^ ~ log *n*. We thus expect that the probability that anisogamy wins will depend primarily on the relative magnitudes of *Ls*^2^ and log *n*, i.e., on the compound parameter *Ls*^2^/ log *n*. This is confirmed by simulations: Fig. 2 shows that for fixed *n* the probability of anisogamous fixation is a function of just *Ls*^2^, while Fig. 3 shows that when *n* is also varied the probability of anisogamous fixation is a function of just the compound parameter *Ls*^2^/log *n*. The above derivation was very rough, ignoring for instance the correction of ≈ 2 log 2, but as our model is a very simplified approximation of any real population, we are just focusing on finding the scaling relationships.

**Figure 2:**
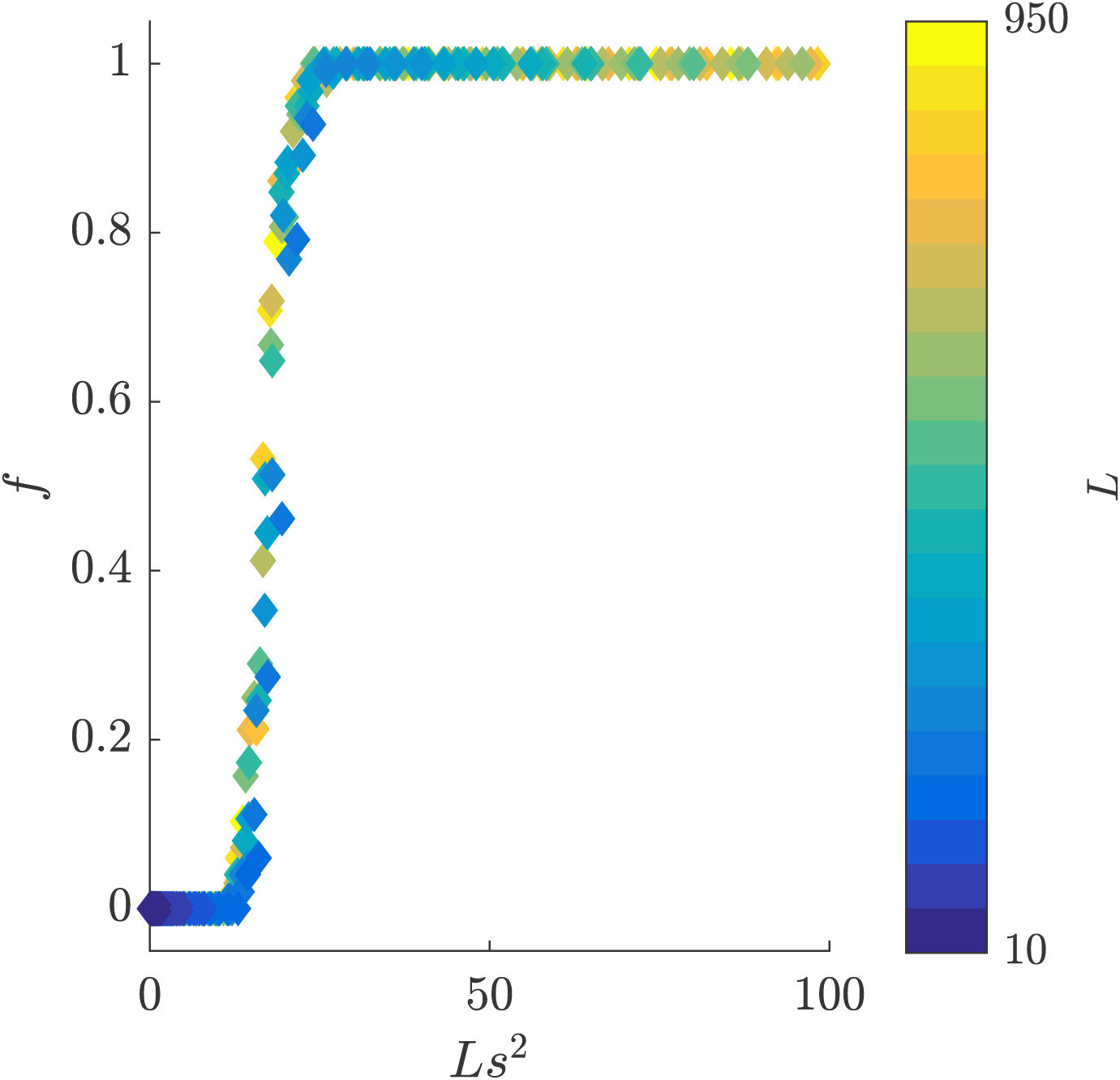
Plot of the frequency of an anisogamous population winning in a competition with the isogamous one over 100 runs, varying *L*, and *s*. Error bars are not shown, but are the binomial error. The color of the markers represents the associated value of *L*. This confirms that the parameter *Ls*^2^ controls the outcome of the competition between the anisogamous and isogamous populations. Parameters: *N* = 1000, *n* = 3.

**Figure 3:**
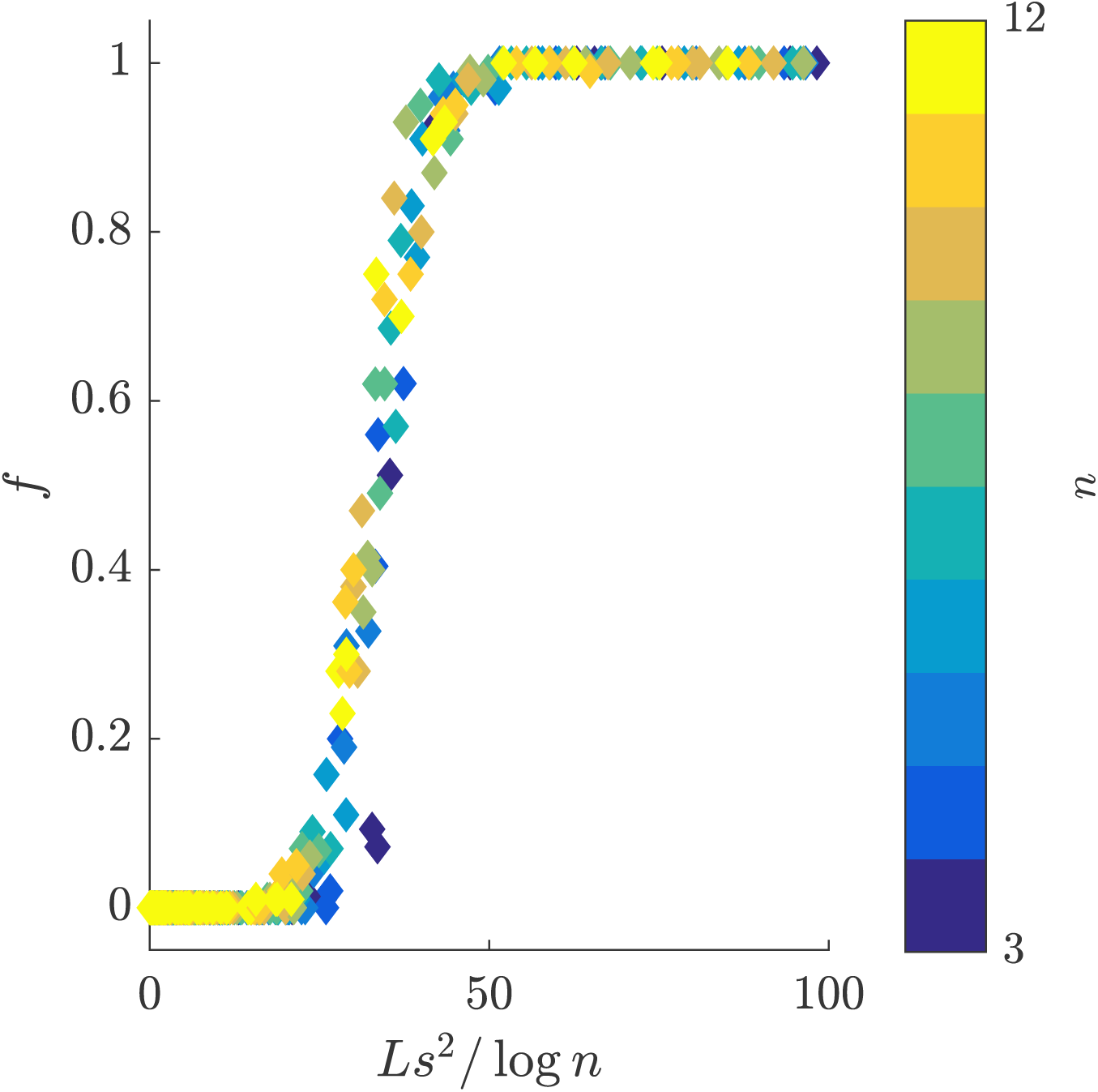
Plot of the frequency of an anisogamous population winning in a competition with the isogamous one over 100 runs, varying *n, L,* and *s*. Like in the previous figure, error bars are not shown, but are the binomial error. The color of the markers represents the associated value of *n*. This confirms that the compound parameter *Ls*^2^/log *n* and not just *Ls*^2^ controls the outcome of the anisogamous/isogamous competition. Here, *N* = 1000.

## 4. Discussion

In this work, we show that the faster spread of beneficial alleles allowed by the presence of males can be sufficient to overcome their two-fold cost and maintain anisogamy in populations adapting to environmental change. By comparing anisogamy to isogamy, we have carefully isolated the effect of sex, rather than confounding it with the effects of recombination. In our minimal model for sexual selection, the key parameter combination determining whether anisogamy is favored over isogamy is the ratio of the variance of potential fitness to the maximal reproductive capacity of isogamous individuals and females, or, equivalently, *Ls*^2^/ log *n*. While we have cast our model in terms of male and female individuals, it would apply equally well to hermaphroditic or monoecious species, or any other mating system as long as each offspring receives resources primarily from one parent.

We have focused on the ability of a fully anisogamous population to out-compete a similarly-sized isogamous population. To fully describe the role of rapid adaptation in the origin and maintenance of anisogamy, two extensions should be considered in the future. First, the assumption that anisogamous and isogamous populations start at the same size should be relaxed. It may be that the mechanism explored here allows the maintenance of anisogamy, but does not allow it to spread when rare. Conversely, it may be that even when the effects discussed here are too weak to be effective under our starting conditions, they can still prevent isogamy from invading an anisogamous population when the isogamous individuals are rare. Second, instead of only considering competition between fully isogamous and fully anisogamous populations, anisogamy should be treated as a quantitative trait, capable of evolving by degrees. A model of competition between alleles that leads to only slightly different degrees of anisogamy could lead to qualitatively different conditions, as carriers of the different alleles would likely interbreed, potentially breaking down the associations between the anisogamy locus and the directly-selected loci [32].

Our model is in some ways very similar to Trivers’ original argument that the two-fold cost of males could be overcome if sexual selection were strong enough so that the average father would be much fitter than the average mother [38]. However, his argument considered only one generation and whether a female could maximize her number of successful daughters via asexual or sexual reproduction, and therefore required potential fathers to be at least twice as fit as mothers for sex to have an advantage. Such a large gap is thought to be rare in natural populations [13]. In our model, we show that because the “good genes” contributed by fathers can compound over time, anisogamy can be favored over the course of multiple generations, even if over a single generation isogamy wins, e.g., in Fig. 1, the anisogamous population is initially increasing. This means that the necessary gap between paternal and maternal fitness can be substantially smaller than in the Trivers’ argument.

Our model does still require quite rapid adaptation for anisogamy to be favored: we must have 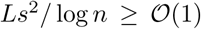, implying variance in fitness of at least 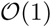. This is more than can be maintained at steady-state by the influx of beneficial mutations [33], although it could be attained temporarily during adaptation to large environmental shifts, as in our simulations. However, this raises the question of what happens when the environment is stable: if anisogamy only has a temporary advantage, with isogamy having a two-fold advantage for long periods before and after, then this mechanism would not seem to be able to contribute substantially to the maintenance of anisogamy. There are two main ways in which this objection can fail.

First, the environment may always be shifting, either constantly exposing new formerly-neutral variation to selection, or fluctuating in direction of the same traits, as is observed on seasonal time scales in *Droshophila* [39]. Selection could also be fluctuating over space, with anisogamous demes successfully adapting to environmental shifts while isogamous ones go extinct. In this sense, our model can also be seen as falling in the general category of Red Queen models for the evolution of sex in continually adapting populations; such models often consider evolving parasites as a source for the environmental shifts [10]. During each environmental shift, genetic hitchhiking will erode the standing neutral variation that could potentially be exposed to selection by future environments: roughly speaking, *N* in Eq. (2) should be replaced by *N_e_*, with log(*N_e_*/*N*) ~ –*Ls*^2^ [33]. But this means that as long as *Nμ*/*n* ≫ 1, i.e., the maternal mutation supply is large, there can still be plenty of genetic variation.

A second possibility is that the threshold rate of adaptation needed for this mechanism to maintain anisogamy may be substantially lower than in our simulations. We have focused on a minimal model of sexual selection, where on a gamete-by-gamete level selection is the same in males and females, with the only difference arising because males have more gametes. But sexual selection can involve much stronger selection on males even on a per-mating basis. If we allow selection on males to be stronger by a factor *α* > 1, this would reduce the threshold for anisogamy to win to 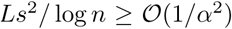, corresponding to a minimum rate of adaptation in females of only 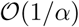. If we imagine, for example, that a mutation allowed dairy cows to reproduce isogamously with each other, this mutation would likely be disfavored, because (artificial) sexual selection is so strong that effectively all selection is taking place in bulls. Note that this can be true even if the rate of increase in the population’s mean milk production, i.e., the rate of adaptation, is not particularly high. On the other hand, in natural populations, such stronger forms of sexual selection might also lead to genetic conflict, in which the alleles that favor male mating success do not improve female fitness, and might also involve direct costs to females in the selection process; these effects would reduce the advantage provided by anisogamy. The experimental evidence for how sexual selection interacts with adaptation to environmental shifts is mixed [24]: some studies have, indeed, found that it accelerates natural selection (e.g., [34]), while others argued that it impedes it (e.g., [40, 41]), or that it can do either depending on the environment [42].

The relevance of our model to the evolution of sex is ultimately an empirical question. If our mechanism is, in fact, a significant contributor, we would expect that anisogamy would typically be more extreme in taxa that have undergone more rapid adaptation in the past. However, this is a very difficult prediction to test. The rate of past adaptation is difficult to measure, particularly so for the fitness flux [43], the relevant quantity here, as opposed to the rate of adaptive substitutions. In addition, the rate of adaptation is likely to correlate with many other factors, all of which could also select for or against anisogamy, confounding the analysis. It may, therefore, make sense to begin by testing our mechanism’s strength in experimental populations. Ideally, this would involve direct competition between individuals differing only in their degree of anisogamy. However, it may be experimentally more tractable to use closely related isogamous and anisogamous species, as are found in, for example, the culturable filamentous fungus genus *Allomyces* [44]. By competing multiple isogamous and anisogamous species and strains against each other under varying degrees of stress, one could test whether the advantage in adaptability conferred by anisogamy is large enough to consistently outweigh the idiosyncratic factors favoring one strain or another. If so, this would be a powerful argument for the potential importance of this mechanism in the evolution of anisogamous sex.

## References

[1] J. Lehtonen, H. Kokko, G. A. Parker, What do isogamous organisms teach us about sex and the two sexes?, Phil. Trans. R. Soc. B 371 (1706) (2016) 20150532.

[2] A. Weismann, The significance of sexual reproduction in the theory of natural selection, Essays upon heredity and kindred biological problems, Clarendon Press, Oxford, 1889.

[3] R. A. Fisher, The genetical theory of natural selection: a complete variorum edition, Oxford University Press, 1930.

[4] H. Muller, Some genetic aspects of sex, The American Naturalist 66 (703) (1932) 118–138.

[5] M. Kimura, T. Maruyama, The mutational load with epistatic gene interactions in fitness, Genetics 54 (6) (1966) 1337.

[6] G. C. Williams, Sex and evolution, Princeton University Press, 1975.

[7] J. Maynard Smith, The evolution of sex, Cambridge Univ Press, 1978.

[8] A. S. Kondrashov, Selection against harmful mutations in large sexual and asexual populations, Genetical research 40 (03) (1982) 325–332.

[9] R. E. Michod, B. R. Levin, The evolution of sex: an examination of current ideas, Sinauer Associates, 1988.

[10] W. D. Hamilton, R. Axelrod, R. Tanese, Sexual reproduction as an adaptation to resist parasites (a review), Proceedings of the National Academy of Sciences 87 (9) (1990) 3566–3573.

[11] N. H. Barton, Why sex and recombination?, Cold Spring Harb Symp Quant Biol 74 (2009) 187–95.

[12] D. Greig, J.-Y. Leu, Natural history of budding yeast, Current Biology 19 (19) (2009) R886–R890.

[13] A. Kodric-Brown, J. H. Brown, Anisogamy, sexual selection, and the evolution and maintenance of sex, Evolutionary Ecology 1 (2) (1987) 95–105.

[14] G. A. Parker, R. Baker, V. Smith, The origin and evolution of gamete dimorphism and the male-female phenomenon, Journal of theoretical biology 36 (3) (1972) 529–553.

[15] F. M. Scudo, The adaptive value of sexual dimorphism: I, anisogamy, Evolution 21 (2) (1967) 285–291.

[16] B. Charlesworth, The population genetics of anisogamy, Journal of theoretical biology 73 (2) (1978) 347–357.

[17] H. Matsuda, P. A. Abrams, Why are equally sized gametes so rare? the instability of isogamy and the cost of anisogamy, Evolutionary Ecology Research 1 (7) (1999) 769–784.

[18] G. Bell, The evolution of anisogamy, Journal of Theoretical Biology 73 (2) (1978) 247–270.

[19] P. A. Cox, J. A. Sethian, Gamete motion, search, and the evolution of anisogamy, oogamy, and chemotaxis, The American Naturalist 125 (1) (1985) 74–101.

[20] R. F. Hoekstra, Why do organisms produce gametes of only two different sizes? Some theoretical aspects of the evolution of anisogamy, Journal of theoretical biology 87 (4) (1980) 785–793.

[21] M. Beekman, B. Nieuwenhuis, D. Ortiz-Barrientos, J. P. Evans, Sexual selection in hermaphrodites, sperm and broadcast spawners, plants and fungi, Phil. Trans. R. Soc. B 371 (1706) (2016) 20150541.

[22] A. Zahavi, Mate selection—a selection for a handicap, Journal of theoretical biology 53 (1) (1975) 205–214.

[23] W. D. Hamilton, M. Zuk, Heritable true fitness and bright birds: a role for parasites?, Science 218 (4570) (1982) 384–387.

[24] U. Candolin, J. Heuschele, Is sexual selection beneficial during adaptation to environmental change?, Trends in ecology & evolution 23 (8) (2008) 446–452.

[25] J. Manning, Males and the advantage of sex, Journal of Theoretical Biology 108 (2) (1984) 215–220.

[26] M. C. Whitlock, Fixation of new alleles and the extinction of small populations: drift load, beneficial alleles, and sexual selection, Evolution 54 (6) (2000) 1855–1861.

[27] A. F. Agrawal, Sexual selection and the maintenance of sexual reproduction, Nature 411 (6838) (2001) 692–695.

[28] S. Siller, Sexual selection and the maintenance of sex, Nature 411 (6838) (2001) 689–692.

[29] L. Hadany, T. Beker, Sexual selection and the evolution of obligatory sex, BMC evolutionary biology 7 (1) (2007) 245.

[30] M. C. Whitlock, A. F. Agrawal, Purging the genome with sexual selection: Reducing mutation load through selection on males, Evolution 63 (3) (2009) 569–582.

[31] D. Roze, S. P. Otto, Differential selection between the sexes and selection for sex, Evolution 66 (2) (2012) 558–574.

[32] M. Kleiman, L. Hadany, The evolution of obligate sex: the roles of sexual selection and recombination, Ecology and Evolution 5 (13) (2015) 2572–2583.

[33] D. B. Weissman, N. H. Barton, Limits to the rate of adaptive substitution in sexual populations, PLoS Genetics 8 (6) (2012) e1002740.

[34] B. Hollis, J. L. Fierst, D. Houle, Sexual selection accelerates the elimination of a deleterious mutant in Drosophila melanogaster, Evolution 63 (2) (2009) 324–333.

[35] G. C. Oakford, Legendary Overbrook Farm stallion Storm Cat dies at 30 (2013). *URL* http://www.drf.com/news/legendary-overbrook-farm-stallion-storm-cat-dies-30

[36] P. D. Lorch, S. Proulx, L. Rowe, T. Day, Condition-dependent sexual selection can accelerate adaptation, Evolutionary Ecology Research 5 (6) (2003) 867–881.

[37] J. F. Crow, M. Kimura, An introduction to population genetics theory, New York, Evanston and London: Harper & Row, 1970.

[38] R. L. Trivers, Sexual selection and resource-accruing abilities in *Anolis garmani*, Evolution 30 (2) (1976) 253.

[39] A. O. Bergland, E. L. Behrman, K. R. O’Brien, P. S. Schmidt, D. A. Petrov, Genomic evidence of rapid and stable adaptive oscillations over seasonal time scales in *Drosophila*, PLoS Genetics 10 (11) (2014) e1004775.

[40] H. D. Rundle, S. F. Chenoweth, M. W. Blows, The roles of natural and sexual selection during adaptation to a novel environment, Evolution 60 (11) (2006) 2218.

[41] S. F. Chenoweth, N. C. Appleton, S. L. Allen, H. D. Rundle, Genomic evidence that sexual selection impedes adaptation to a novel environment, Current biology 25 (14) (2015) 1860–1866.

[42] L. Yun, P. J. Chen, A. Singh, A. F. Agrawal, H. D. Rundle, The physical environment mediates male harm and its effect on selection in females, Proceedings of the Royal Society B: Biological Sciences 284 (1858) (2017) 20170424.

[43] V. Mustonen, M. Lässig, Fitness flux and ubiquity of adaptive evolution, Proceedings of the National Academy of Sciences 107 (9) (2010) 4248–4253.

[44] S. Phadke, S. Rupp, M. Wilson Sayres, Understanding the evolution of anisogamy in the early diverging fungus, Allomyces, bioRxivdoi: 10.1101/230292.

